# WHATEVER YOU WANT: INCONSISTENT RESULTS IS THE RULE, NOT THE EXCEPTION, IN THE STUDY OF PRIMATE BRAIN EVOLUTION

**DOI:** 10.1101/454132

**Authors:** Andreas Wartel, Patrik Lindenfors, Johan Lind

## Abstract

Primate brains differ in size and architecture. Hypotheses to explain this variation are numerous and many tests have been carried out. However, after body size has been accounted for there is little left to explain. The proposed explanatory variables for the residual variation are many and covary, both with each other and with body size. Further, the data sets used in analyses have been small, especially in light of the many proposed predictors. Here we report the complete list of models that results from exhaustively combining six commonly used predictors of brain and neocortex size. This provides an overview of how the output from standard statistical analyses changes when the inclusion of different predictors is altered. By using both the most commonly tested brain data set and a new, larger data set, we show that the choice of included variables fundamentally changes the conclusions as to what drives primate brain evolution. Our analyses thus reveal why studies have had troubles replicating earlier results and instead have come to such different conclusions. Although our results are somewhat disheartening, they highlight the importance of scientific rigor when trying to answer difficult questions. It is our position that there is currently no empirical justification to highlight any particular hypotheses, of those adaptive hypotheses we have examined here, as the main determinant of primate brain evolution.

## Introduction

The field of primate brain evolution can be characterized as an array of contradicting results (1, 2). Most studies have utilized phylogenetic comparative methods, in the guise of phylogenetic generalized least squares regression (PGLS). Brain or neocortex size have often been the dependent variables, in combination with a varying number of predictor variables, depending on the hypothesis at hand. As conflicting results abound we think an evaluation of this approach has its merits. Therefore, we here systematically vary choice of data set and inclusion/exclusion of predictor variables in the PGLS framework, to investigate if, why, and when contradictory results emerge.

Most previous studies have relied on one of only two available datasets on brain size (3, 4). We here include new data (5) added to one of the old datasets (3) to broaden the reanalysis. Though eager to reach interesting biological conclusions from the new data, we are foremost concerned with evaluating the validity of previous analyses. We believe that if the data is underdetermined in the sense that predictor A is significant and predictor B non-significant in one context, while the reverse is true in another context, then we have currently no method to determine which variable is most important, if any. As will be explained, our choices of both method and data are based on what is praxis in the field of primate brain evolution – this study is not driven by any particular biological hypothesis and seeks only to reach biological conclusions about reliability of results.

There exist many suggested non-mutually exclusive hypotheses for causes of variation in size and architecture of primate brains. Here, we summarize seven such particularly popular hypotheses that have been both supported and rejected in various studies.

### Allometric relationships

Brains are similar to other organs and thus scale allometrically with body size. Similarly, brain parts scale allometrically with brain size. Simply put, larger brains are required to run larger bodies. Most differences in brain size and brain architecture can thus be predicted by body size (6, 7)). Due to such known allometric relationships, one usually controls for body size / brain size in studies of primate brains. Whatever residual variation is left is the target for the adaptive hypotheses. The rationale here is that “intelligence” corresponds to the amount of excess brain mass after controlling for brain mass dedicated to bodily functions (8, 10, 11, 12, but see 9). However, body size alone accounts for more than 90% of the variation in brain size differences between primates (13, 8), so only little is left to explain.

### General cognitive abilities

Larger relative brain size or brain component size has evolved to meet higher cognitive demands (8, 14, 15, 12, 16, 17, 18).

### The social brain hypothesis

Some primates evolved large brains and/or larger brain components for reasons having to do with social complexity (e.g. 19, 20, 21, 22, 23, 24, 25).

### Sexual selection

Demands of sociality are different between males and females. This should produce detectable differences in relative brain size or brain component size between species where sexual selection is high compared to species where it is relaxed (26, 27, 10, 28).

### Diet

Fruits are harder to find than leaves, so some primates have evolved larger brains and/or large specific brain parts to cope with “a challenging diet of fruit” (29 p. 312, 30, 31, 32, 33). Alternatively, the causal relationship is hypothesized to go in the other direction, that larger brains (that evolved for another reason) demands a more high-calorie diet (34, 35).

### Life history

Variation in juvenile period and life span is hypothesized to affect brain size evolution (36, 37). An extended juvenile learning is necessary to evolve a bigger brain (38, 39). Also, a longer life span is a consequence of slow growth in order to cope with the high energy costs of developing a large brain (40, 41, 42) and/or to facilitate more opportunities to harness the products of enhanced brain size (36, 43).

### The mosaic brain hypothesis

This is a composite hypothesis where it is hypothesized that “variation in the size of individual brain components reflects adaptive divergence in brain function mediated by selection” (44, p. 2, 45, 46, 27). Here, all hypotheses can come into play simultaneously (21).

The list of competing hypotheses can go on (1, 36, 2), but the message from the literature is clear: there is no real consensus about the adaptive explanations for neither primate brain size nor primate brain architecture. All these studies have sought to find evolutionary drivers of primate brain evolution, where residual brain size or different aspects of brain architecture have been used as approximations of intelligence, making it difficult and unjustified to highlight any particular hypothesis from the smorgasbord of significant results.

Because results have proven both ambiguous and contradictory, we use new data (5) in combination with the classic brain data set provided by (3) and report the complete list of models that results from exhaustively combining six commonly used predictors (female group size, male group size, female sexual maturity, life span, innovation, and percent fruit in diet). Our choice of predictors in this studie reflect our notion of what hypotheses is common and is not an exhaustive combination of tested predictors. Others have used combinations of other predictors (e.g. 37, 47).

We start out by calculating the ‘best’ model according to the Akaike information criteria (AIC) both when using total brain size as the dependent variable and when using neocortex size as the dependent variable. Then we use this output and examine the stability of results when the inclusion of different predictors is altered. We end by examining the stability of previously published analyses in the same way.

Our aim here is not to reach a final verdict on the biological relevance of different hypotheses, but rather to investigate if data and methods currently at hand are productive enough for such considerations in the first place.

## Material & Methods

### Data

All data used in this study were collected from published literature and are presented in the Appendix. Most studies on primate brain evolution have relied on a classic data set on primate brains provided (3), for example (31, 32, 48, 49, 41, 36, 25, 50, 19, 35, 47, 38, 51, 52, 53, 54, 55, 26, 27, 56, 44, 57, 58, 59, 14, 60, 20, 61, 62, 63, 39, 64). Data on brain and neocortex size used in this study were obtained by pooling (3) and new data from (5). By pooling we refer to the weighted average of two or more sets of data.

Life history and body size data were obtained by pooling data from (65) and (33). Length of juvenile period is approximated by age of sexual maturity. Life span is calculated as the period between sexual maturity and maximum recorded age at death. Percentage fruit in diet were obtained by pooling data from (33, 66, 65, 67). Rates of innovation were taken from (14).

We used female weight as a proxy for species weight as it is less variable than male weight among species. Variation in male weight is in sexually selected species to a large degree a consequence of selection on physical strength (68).

Though there are several ways to quantify social complexity, e.g. pair-bonding (20), tactical deception (25), we use group size as it is the most common used approximation of social complexity (e.g. 69, 51, 26, 27, 54, 38, 30, 39). In this study both male and female group sizes are used because it has previously been shown that female rather than male group size correlate positively with neocortex volume in primates (26; 27), suggesting that it is social demands on females that mainly drives primate brain evolution.

Number of species with full data on all variables (including phylogeny) and included in analyses are *N = 40.*

### Statistical analysis

All analyses were executed in R (70) using the packages NLME (71), APE (72), MASS (73) and BRMS (74). All variables were log-transformed prior to analysis except percentage fruit in diet, which instead were arcsine-square root transformed.

We used phylogenetic generalized least squares (PGLS) regressions throughout. This method allows for the estimation of the impact of phylogeny on the covariance among residuals, thereby controlling for relatedness (75, 76). A consensus phylogeny for each dataset were obtained from (77). Lambda (*λ*) was estimated but in some cases when lambda is very close to 1, processing in R sometimes crash due to an optimization error. When this happened lambda was fixed to 1 (77).

All combinations of the following variables were used as predictors: female weight, female group size, male group size, female sexual maturity, life span, innovation, and percent fruit in diet, both when using total brain size and neocortex as the outcome. The mosaic brain hypothesis (see introduction) is explicitly tested when using neocortex as the outcome variable. Female weight was included as independent variable in all analysis (i.e. in 63 models) because it is standard procedure to control for body and thereby consider the analyses as predicting *relative brain size* (but see for example 12). This sums to 63 models per dependent variable (total brain and neocortex).

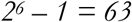

It therefore follows that each predictor is included in 32 * 2 models.

Model selection was carried out utilizing the Akaike information criteria (AIC).

One way to assess the severity of collinearity in a least squares regression is to calculate the variance inflation factor (VIF). However, because the PGLS regression used here assumes a correlated residual structure the VIF diagnostic does not carry over easily (75). Our approach was instead to calculate posterior distributions in a Bayesian framework and visually inspect weather they correlate (Appendix).

## Results

We analyzed the effect of six predictor variables on two outcome variables: total brain size and neocortex size. First, we calculated the ‘best’ model according to the Akaike information criteria (AIC), both when using total brain size as the dependent variable and when using neocortex size as the dependent variable. As can be seen in table 1, AIC resulted in a model that includes female weight, male group size, female group size, lifespan, female sexual maturity and fruit, omitting only innovation, as the best model predicting total brain size. Likewise, in table 2 for neocortex size as the dependent variable, AIC resulted in a model that includes female weight, male group size, female group size and female sexual maturity.

**Table 1.** The following model was selected with AIC for total brain size as the dependent variable.

| *Predictor* | *B* | *se* | *t* | *p* |
| --- | --- | --- | --- | --- |
| *Female weight* | *0.590* | *0.039* | *15.177* | *<0.000* |
| *Male group size* | *-0.090* | *0.058* | *-1.551* | *0.131* |
| *Female group size* | *0.108* | *0.049* | *2.220* | *0.033* |
| *Life span* | *0.240* | *0.086* | *2.800* | *0.009* |
| *Female sexual maturity* | *0.158* | *0.090* | *1.756* | *0.088* |
| *Fruit* | *0.238* | *0.088* | *2.720* | *0.010* |
| *Model summary* |  |  |  |  |
| *R<sup>2</sup>* | *0.975* |  |  |  |
| *λ* | *1* |  |  |  |
| *N* | *40* |  |  |  |

**Table 2.**
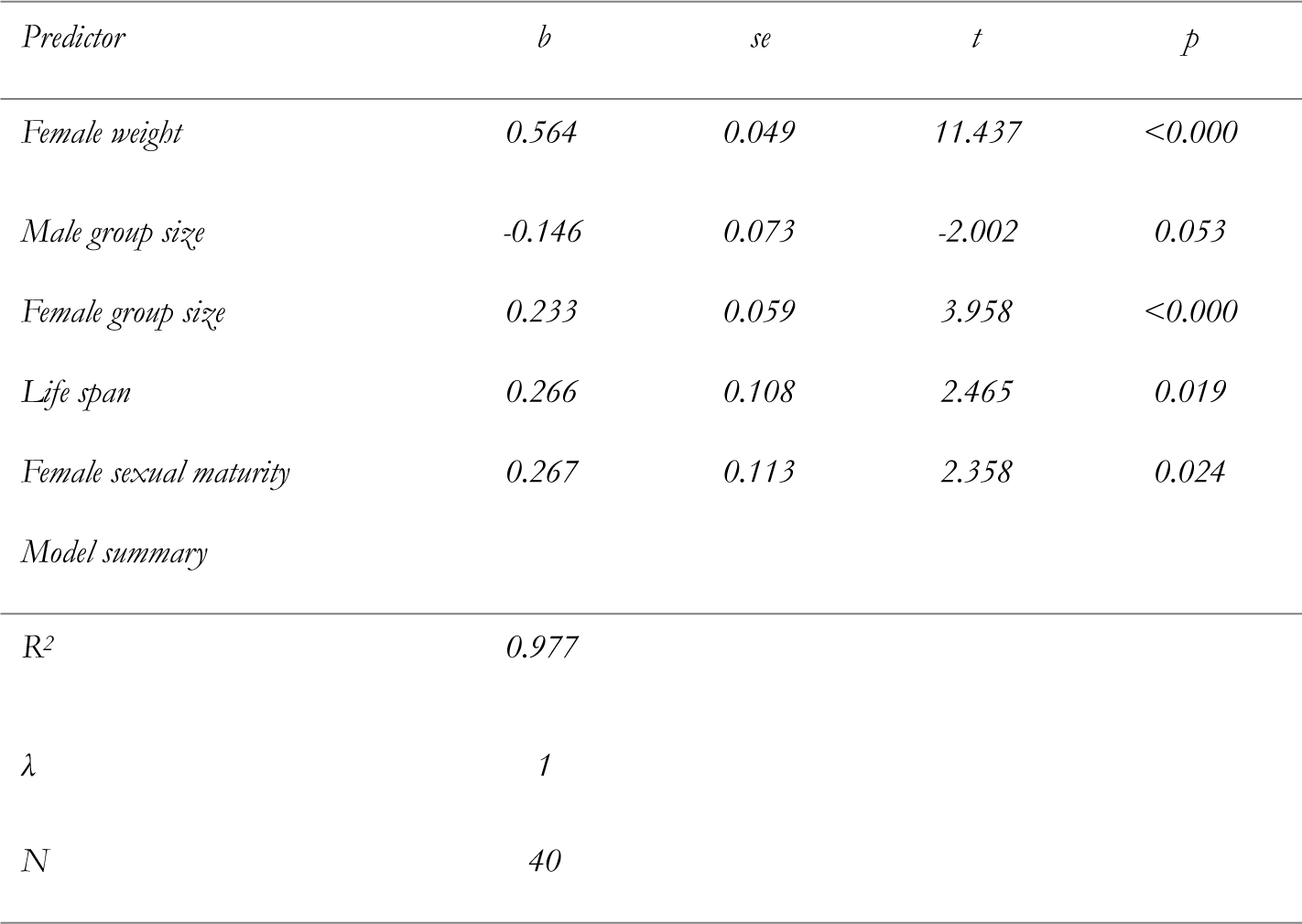
The following model was selected with AIC for neocortex size as the dependent variable.

| Predictor | *b* | *se* | *t* | *p* |
| --- | --- | --- | --- | --- |
| Female weight | 0.564 | 0.049 | 11.437 | <0.000 |
| Male group size | -0.146 | 0.073 | -2.002 | 0.053 |
| Female group size | 0.233 | 0.059 | 3.958 | <0.000 |
| Life span | 0.266 | 0.108 | 2.465 | 0.019 |
| Female sexual maturity | 0.267 | 0.113 | 2.358 | 0.024 |
| Model summary |  |  |  |  |
| R<sup>2</sup> | 0.977 |  |  |  |
| $\lambda$ | 1 | | | |
| N | 40 |  |  |  |

AIC is not a method that involves p-values *per se*. Instead it estimates the out-of-sample deviance and is therefore concerned with prediction. However, in the literature of primate brain evolution, p-values are often used to select among AIC-selected predictors with the aim to reinforce or undermine hypotheses. Therefore, we calculated p-values for the six predictors for all models possible, that is the exhaustive combinations of the six predictors. This resulted in 63 models in total where each predictor was included 32 times. Many predictors were estimated both above and below *p* = 0.05, the conventional boundary for the rejection of hypotheses. Whether a predictor was above or below *p=0.05* depended on what other variables were included in a particular model (i.e. which concomitant predictors). Table 3 and 4 illustrate this by first showing the model in which each predictor was estimated to the lowest p-value, and subsequently the model that gave the highest p-value. As can be seen in table 3, where total brain was used as the dependent variable, female group size, life span, female sexual maturity and fruit each moved from significant to non-significant when changing concomitant predictors. Likewise, as shown in table 4 where neocortex was used as the dependent variable, male group size, life span and female sexual maturity each moved from significant to non-significant when changing concomitant predictors.

**Table 3.** *Shows the change in p-value for each predictor when altering concomitant predictors using total brain as dependent variable. Read as follows: the focal predictor in the first column was estimated to a lowest p-value (out of all the 32 models the focal predictor where included) shown in the second column when using concomitant predictors shown in column three. Likewise, the maximum p-value shown in column four, were estimated using concomitant predictors in column five. N = 40.*

| *Focal predictor* | *Min p-value* | *Concomitant predictors* | *Max p-value* | *C o n c o m i t a n t*<br>*predictors* |
| --- | --- | --- | --- | --- |
| *Male group size* | *0.082* | *Female group size, Lifespan, Female sexual maturity, Innovation, Fruit* | *0.968* | *Lifespan* |
| *Female group size* | ***0.017*** | *Male group size, Lifespan, Female sexual maturity, Innovation, Fruit* | *0.540* | *Female sexual maturity, Innovation* |
| *Life span* | ***0.004*** | *Male group size, Female group size, Female sexual maturity, Innovation, Fruit* | *0.083* | *Male group size, Innovation* |
| *Female sexual maturity* | ***0.043*** | *Male group size, Female group size, Life span, Innovation* | *0.263* | *Female group size, Innovation, Fruit* |
| *Innovation* | *0.240* | *Fruit* | *0.865* | *Male group size, Life span* |
| *Fruit* | ***0.007*** | *Male group size, Female group size, Lifespan* | *0.061* | *Male group size, Female sexual maturity, Fruit* |

**Table 4.** *Shows the change in p-value for each predictor when altering concomitant predictors using neocortex size as dependent variable. Read as follows: the focal predictor in the first column was estimated to a lowest p-value (out of all the 32 models the focal predictor where included) shown in the second column when using the concomitant predictors shown in column three. Likewise, the maximum p-value shown in column four, were estimated using concomitant predictors in column five. N = 40*

| *Focal predictor* | *Min p-value* | *Concomitant predictors* | *Max p-value* | *C o n c o m i t a n t*<br>*predictors* |
| --- | --- | --- | --- | --- |
| *Male group size* | **0.039** | *Female group size, Lifespan, Female sexual maturity, Innovation* | *0.780* | *Lifespan, Female sexual maturity, Innovation, Fruit* |
| *Female group size* | **0.0003** | *Male group size, Lifespan, Female sexual maturity* | **0.015** | *Female group size, Female sexual maturity, Innovation, Fruit* |
| *Life span* | **0.017** | *Male group size, Female group size, Female sexual maturity, Innovation, Fruit* | *0.403* | *Male group size, Innovation* |
| *Female sexual maturity* | **0.016** | *Male group size, Female group size, Life span, Innovation, Fruit* | *0.153* | *Male group size, Innovation* |
| *Innovation* | *0.074* | *No concomitant predictors* | *0.970* | *Male group size, Female group size, Life span* |
| *Fruit* | *0.138* | *Lifespan, Female sexual maturity, Innovation* | *0.699* | *Male group size, Female group size, Lifespan, Innovation* |

To get an overview of all models, i.e. the 32 models that each predictor were included in, table 5 and 6 illustrate the number of models in which each predictor was non-significant. Table 5 shows models that used total brain as the dependent variable, whereas Table 6 shows the same models using neocortex size as the dependent variable. As can be seen, whether a variable is a significant predictor of brain or neocortex size depends to a worryingly high degree upon what concomitant variables that were also included in the model.

**Table 5.** *Number of models in which each predictor was estimated non-significant (p > 0.05) with total brain size as the dependent variable. N = 40*

| *Predictor* | *Number of models in which predictor is non-significant* |
| --- | --- |
| *Female weight* | *0/32* |
| *Male group size* | *30/32* |
| *Female group size* | *28/32* |
| *Life span* | *6/32* |
| *Female sexual maturity* | *30/32* |
| *Innovation* | *32/32* |
| *Fruit* | *6/32* |

**Table 6.** *Number of models in which each predictor were estimated non-significant (p > 0.05) with Neocortex size as the dependent variable. N = 40.*

| Predictor | Number of models in which predictor is non-significant |
| --- | --- |
| Female weight | 0/32 |
| Male group size | 32/32 |
| Female group size | 0/32 |
| Life span | 26/32 |
| Female sexual maturity | 21/32 |
| Innovation | 32/32 |
| Fruit | 32/32 |

To further illustrate how predictors jump above and below the significance level *p* = 0.05, table 7 shows results that extend beyond the analyses hitherto reported. Here we have reanalyzed previously reported results by systematically altering one factor at a time and observe changes in calculated p-values. The first row in table 7 shows that the significant relationship reported by (33), between fruit in diet as the predictor and brain size as the dependent variable, becomes non-significant when updating all predictors, that is adding data by pooling. Row two shows the opposite change. (33) reported a non-significant relationship between group size and brain size. However, with other brain data (i.e. changing data from [4] to [13] & [5]) the relationship is significant. Row three shows that (38) reported relationship between juvenile period and brain size reverse from significant to non-significant when adding more data to predictors by pooling. Row four shows that the relationship reported by (27) between various brain parts and sexual dimorphism, female group size and male group size reverse or disappear when, again, adding more data by pooling. Lastly on row five, (19) reported a significant relationship between group size and neocortex with a slope that significantly changed when adding more data by pooling and controlling for phylogeny. Further, ‘Dunbar’s number’, claimed to describe the cognitive threshold for group size in humans, changed from 150 to 22. Note also that the 95% confidence interval for this number ranges from 0.000002 to 251,549,413, rendering the threshold number useless (the asymmetry of the confidence interval stems from exponentiating the fitted log[Y]). Note that when predicting with PGLS the model does not account for the phylogenetic position of the observation to be predicted. All reanalyzes reported here use phylogeny to correct for non-independence.

**Table 7.** Overview of changes in the relation between brain size and predictors as different data is used.

| *Source* | *Reported relationship* |  | *Reanalysis changes* | *Result* |
| --- | --- | --- | --- | --- |
| (33) | Brain size ~ Fruit in diet | Significant | Updated predictors<sup>1</sup> | Non-significant |
| (33) | Brain size ~ Group size | Non-significant | Other brain data<sup>2</sup> | Significant |
| (38) | Juvenile period ~ non visual neocortex | Significant | Updated variables<sup>3</sup> | Non-significant |
| (27) | Specific brain parts ~ Dimorphism, Female/male group size | Non-significant/<br>Significant | Updated variables<sup>4</sup> | Significant/<br>Non-significant |
| (19) | ‘Dunbar’s number’ = 150 |  | Updated variables and control for phylogeny<sup>5</sup> | ‘Dunbar’s number’ = 22 |

<sup>1</sup>Using (33) brain data but pooling predictors from several sources (see Appendix). <sup>2</sup>Using predictors from (33) but changing their brain data (4) to this study’s data (i.e. pooling [3, 5]). <sup>3</sup>All variables used were pooled with data from (38, 65, 33). <sup>4</sup>All variables used were pooled with data from (65, 33). <sup>5</sup>For brain size data, we added the new data from (5) to that of (3). Further, data on group size and body weight was pooled from (65, 33). Dunbar’s original model was used to predict *Homo sapiens* group size (‘Dunbar’s number’), without control for phylogeny. Note that in addition to being very different from the original estimate, the new ‘Dunbar number’ from our more complete and phylogenetically controlled model has a huge 95% confidence interval, ranging from 0.000002 to 251,549,413. For all results see the Appendix.

As is custom in phylogenetic comparative analysis, phylogenetic information is used to estimate the covariance of the residuals (78). This process can lead to an *R*^2^ value different from model fit with non-phylogenetic least squares. With this in mind it can still give a crude picture of the amount of brain size variation that is explained by body size: *R*^2^ = cor(predicted, log of total brain size)^2^ = 0.94, where predicted(total brain) = *intercept*(5.167) + *b*(0.667)*log of female weight.

## Discussion

Our analyses indicate that the field of primate brain evolution is best characterized as an array of contradicting results (1, 2) and our results reveal why this is so. Within the PGLS framework, choice of what variables to include, and what observations for those variables to include, fundamentally changes the conclusions as to what drives primate brain evolution. In this study we included new data (5) with the classic dataset on primate brain size (3) but this did not change the volatility in the results. If so inclined, we could have presented support for any pet-hypothesis of our choice, and refuted any study we would have liked. Combined with the ‘publish-or-perish’-situation in academia, this is not an ideal situation.

In table 1 and 2 we present the models that were selected with AIC. The AIC test in turn had six explanatory variables to combine. The predictors and the AIC test itself were chosen according to our best effort to follow the established method within the field of primate brain evolution. In other words, we chose variables that according to the literature are plausible determinants of brain size, and we used state-of-the-art methods to choose among combinations of predictors. If this had been a standard study, we would have moved on to discuss the biological rationale for these AIC chosen models and special attention would have been given to the significant predictors (at *p* < 0.05). However, we argue that because of the breadths of hypotheses or hierarchies of hypotheses, compatible with the results, the more important aspect of this study is the instability of results (tables 3-7).

Tables 3-6 show that most of the explanatory variables have been assigned parameter values with probability on both sides of the significance level at *p* = .05. Table 3 and 4 show the most extreme cases in the exhaustive list of models. Thus, using p-values to evaluate the importance of hypotheses that affect primate brain size leaves us ambivalent. AIC was developed to select among models and thus to save us from such ambivalence, but AIC can only evaluate the models given to it, which is why the results still are dependent on pre-test variable choice.

Even though AIC has been established as praxis, many papers on primate brain evolution gives special status to predictors associated with *p* < .05. Following this habit, we think there is no way avoiding the problem illustrated in table 3 - 6; depending on what initial predictors happened to be included in the analyses, they can either be judged important (*p* < 0.05) or non-important (*p* > 0.05) by the order of magnitude shown in table 5 and 6 (for the predictors included in this study). The mosaic brain hypothesis - explicitly tested when using neocortex as the outcome variable - did not escape the problem of inconsistent result as can be shown in table 4 and 6.

When we included new brain data and updated variables on previously reported results, we found the same patterns. As shown in table 7, the results reported by (33) indicated that brain size was best predicted by diet, but not by sociality (measured as group size). When we added more observations (see Appendix) to their explanatory variables, both diet and sociality turned out to be non-significant. When we kept their original predictors, but used pooled brain data (3, 5), sociality became significant but not diet (see Appendix).

Further, (27) predicted that the relative size of brain structures involved in motor skills and coordination, such as the mesencephalon, diencephalon, cerebellum and medulla oblongata, would increase in species with greater rates of sexual size dimorphism. They found dimorphism to be a significant predictor for all these structures except cerebellum. However, when we did a reanalysis with updated variables we found no significant relationships for any of these predictors.

These results confirm a previous report: (1) re-analyzed endocranial data provided in (4) and concluded that “[o]ur results indicate that, even holding constant statistical methods, phylogeny, set of predictor variables, response variable data, and species sample, the behavioral and ecological correlates of brain size are sensitive to the use of different predictor datasets” (1, p. 4).

AIC is a method for choosing the model with the lowest out-of-sample deviance and as such a method concerned with prediction, not p-values. Clearly, as shown in this paper, the best predicting model may include variables that have non-significant p-values. In the context of AIC, it is easy to illustrate that the best predicting models sometimes do not reveal the true relationship between individual predictors and the outcome, as for example in the case of concomitant variable bias (79) or collinearity. Yet inference about individual predictors is mostly what concerns scientists of primate brain evolution, not mere prediction.

Collinearity is member of a family of problems with model fitting referred to as weakly-identifiable parameters (or sometimes non-identifiable) (80). If the predictors co-vary a lot, i.e. share information, their posterior parameter distributions will correlate (when *β*_1_ increase *β*_2_ must decrease and vice versa) making it hard to identify a true estimate. To investigate if our analyses suffer from collinearity we calculated posterior distributions for all parameters in a Bayesian framework, for the full model (containing all six predictors) and plotted the correlation matrix (excluding varying intercepts, see Appendix). We conclude from visual inspection that for some parameters there exist substantial correlation that could explain the varying results exposed in this study (80, 81, 82).

Further caveats have been raised by other researchers, such as problems with measuring and comparing intelligence (15; 83), the idea of adaptive specializations of cognitive mechanisms (84, 85), validity of observational data versus experiment (18), choice of brain measure (50, 6), measuring and defining sociality (86, 87, 69) and p-hacking: given that the same sample on brain volume (3) has been modelled against many variables, it is to be expected that *Type 1*-errors will emerge (88). Also, there is some evidence that different data samples are qualitatively different from each other (89, 90, 69). It has for example been shown that data on body size often are averaged, inaccurate and from unspecified sources (91, 89).

In our analyses, variation in body size explain 94% of the variation in brain size. This does not leave much to be explained by the competing adaptive hypotheses. The fact that these adaptive hypotheses explain very little variation, taken together with the unstable nature of results, suggest that it is easy to overstate the importance of sociality, diet, problem solving, or life-history for understanding brain evolution.

Further, other measures not included here may be more important for our understanding of brain evolution. Indeed, other combinations of predictors have been used in previous studies, however, we believe that adding more predictors would reveal similar inconsistencies in the results and that the six predictors used in this study suffice to illustrate this. That variation in sensory and perceptual systems give rise to variation in brain size is not controversial (92, 93). A primate with very large eyes will have brain areas that correspond to sensory and perceptual needs. In addition, animals that are motor flexible, have many different kinds of muscles, and large behavior repertoires need brain areas that control muscles. Therefore, larger brains are needed to drive more motor flexible bodies (94). To put this in Tinbergian terminology: a mechanistic link between brain size and body functions is straight forward and non-controversial, while a functional link between brain size and mental capacities is harder to define to non-controversial precision.

It is our position that, given the data at hand and the PGLS approach, there is currently no empirical justification to highlight any particular hypotheses, of those adaptive hypotheses we have examined here, as the main determinant of primate brain evolution.

## Acknowledgements

This research project was supported by Knut and Alice Wallenberg Foundation Grant 2015.0005 (JL) and Marianne and Marcus Wallenberg Foundation Grant 2017.0049 (PL). We are grateful for input from two anonymous reviewers that greatly improved the manuscript.

## Data Availability

All data is available in the Appendix.

## Author Contribution

AW, PL, JL contributed equally, although AW performed the statistical analyses.

## Research Ethics

No ethical assessment were needed because the study used published data.

## Animal Ethics

This study was performed without any animal subjects and no approval from ethics committees was needed.

